# iSsus3744: A Genome-Scale Model-Guided Strategy for Rational Media Design for Cultivated Pork

**DOI:** 10.64898/2026.05.28.728221

**Authors:** Sandra Gomez Romero, Mark Vigliotti, Victoria Ramirez Lopez, Kyle Nguyen, Valeria Marchitto, Nanette Boyle

**Affiliations:** Quantitative Biosciences and Engineering, Colorado School of Mines, Golden, CO, USA; Chemical and Biological Engineering, Colorado School of Mines, Golden, CO, USA; Cultivated Meat Modeling Consortium

**Keywords:** cultivated meat, genome scale metabolic model, metabolic flux analysis, media optimization, bioprocess

## Abstract

Cultivated meat production is currently limited by high production costs and an incomplete understanding of cellular metabolism in agriculturally relevant species. Genome-scale metabolic models (GEMs) have successfully guided media optimization in biopharmaceutical systems but have not been widely applied to cultivated meat. In this study, we present iSsus3744, the first genome-scale metabolic reconstruction for *Sus scrofa* and demonstrate its application for rational media design in cultivated pork production. iSsus3744 was reconstructed using HumanGEM and Recon3D as template models and further constrained using experimentally determined biomass composition and uptake and excretion fluxes from a Duroc porcine muscle satellite cell line. The final model comprised 3,744 genes, 8,854 metabolites, and 12,248 reactions distributed across eight cellular compartments. Flux balance analysis (FBA) and flux variability analysis (FVA) were used to identify amino acids limiting cellular growth and predict media supplementation strategies. Experimental validation demonstrated that model-guided amino acid supplementation significantly improved proliferation. Supplementation with phenylalanine reduced doubling time from 31.9 hours to 17.2 hours, representing a 46% reduction, while lysine, methionine, tyrosine, leucine, and valine also improved growth performance. These results demonstrate the potential of genome-scale metabolic modeling as a powerful platform for rational media optimization in cultivated meat systems. iSsus3744 provides a foundational resource for future integration of omics, transcriptional regulation, and isotope-assisted metabolic flux analysis to further accelerate serum-free media development and cultivated meat bioprocess optimization.

## INTRODUCTION

The field of cellular agriculture integrates stem cell biology, tissue engineering, biomaterials, bioprocessing, and animal science to produce animal-derived products in vitro (O’Neill et al., 2021; Reiss et al., 2021). The field draws on expertise in engineering, biology, sustainability, and economics to establish robust cell lines, develop efficient cell culture media, and design edible biomaterials and plant-based serum-free media formulations (Block, 2020). Despite rapid advances in the field, cell culture media remains one of the largest economic barriers to large-scale production, accounting for approximately 33% to 90% (Humbird, 2021; Specht, 2020) of overall manufacturing costs. As a result, improving and accelerating media optimization has become a major focus of research in cellular agriculture. Approaches such as artificial intelligence (Nikkhah et al., 2023), spent media analysis (O’Neill et al., 2022), and systems biology-based modeling (Gomez Romero & Boyle, 2023) have all been proposed as strategies to reduce development time and improve media performance.

Among these approaches, genome-scale metabolic models (GEMs) provide a powerful framework for understanding and predicting cellular metabolism. GEMs are network-based reconstructions that integrate genes, metabolites, metabolic pathways, biochemical reactions, and Gene-Protein-Reaction (GPR) relationships to describe the metabolic capabilities of a cell or organism. These models can be further refined through the integration of multi-omics datasets, including genomics, transcriptomics, epigenomics, proteomics, and metabolomics, enabling more biologically accurate and condition-specific predictions. The integration of omics-constrained GEMs has also been proposed as a valuable platform for machine learning applications in biotechnology (Passi et al., 2021). GEMs have already demonstrated significant utility in biotechnology and food production. In the food industry, they are used to better understand microbial metabolism in fermentation processes, with applications in improving product flavor, optimizing fermentation efficiency, and enhancing food quality and safety (Somerville et al., 2022). Similarly, GEMs have been successfully applied to optimize cell culture media in mammalian bioprocessing systems. In 2020, Huang et al. used the Hefzi et al. 2016 Chinese Hamster Ovary (CHO) genome-scale model (Hefzi et al., 2016) to design improved media formulations for CHO-K1 cells (Huang et al., 2020). The authors first added cell line specific constraints based on RNA sequencing data collected during exponential and stationary growth phases, mapping 1,728 upregulated genes onto the model. Experimentally measured nutrient uptake and metabolite secretion rates were then incorporated as metabolic constraints. Using Flux Balance Analysis (FBA), the authors predicted that increasing leucine and valine concentrations would enhance IgG productivity by 46%. Experimental validation confirmed these predictions, resulting in a 33% increase in IgG production in the optimized media (Huang et al., 2020). These studies demonstrate the potential of GEM-guided approaches for rational media optimization.

Although pork is the third largest global source of animal protein after fish and chicken, a comprehensive genome-scale metabolic reconstruction for pigs has not previously been available. Beyond their agricultural importance, pigs are also highly valuable biomedical model organisms because of their strong anatomical and physiological similarities to humans (Lunney et al., 2021). We have previously proposed the use of genome-scale metabolic modeling as a strategy to optimize cell culture media for cultivated meat applications (15, 16). In addition, we reviewed the current state of metabolic modeling for agriculturally relevant species, including cattle (*Bos taurus*), pigs (*Sus scrofa*), chicken (*Gallus gallus*), salmon (*Salmo salar*), shrimp (*Penaeus* spp. and *Litopenaeus* spp.), and duck (*Anas platyrhynchos domesticus*) (15), identifying major gaps in the availability of GEMs for cultivated meat research. Here, we address one of these gaps by presenting the first genome-scale metabolic reconstruction for *Sus scrofa*, named iSsus3744. The model was constrained using experimentally determined biomass compositions and nutrient uptake and excretion rates. We then used iSsus3744 to identify amino acids limiting cell growth and proliferation and to predict media compositions that improve cellular performance.

## MATERIALS AND METHODS

### Model reconstruction

The RAVEN (Reconstruction, Analysis and Visualization of Metabolic Networks) toolbox (version 2.8.4), and the COBRA Toolbox (version 2.34.1) (Ebrahim et al., 2013; Heirendt et al., 2019; Schellenberger et al., 2011; Vlassis et al., 2014) were used to construct a draft ortholog model (Agren et al., 2013; Wang et al., 2018). This draft ortholog model was created using HumanGEM v1.6.0 (Robinson et al., 2020) and Recon3D (Brunk et al., 2018) as a template model. Next, the Kyoto Encyclopedia of Genes and Genomes (KEGG) (Kanehisa et al., 2023; Kanehisa & Goto, 2000; Kanehisa, 2019) was used to create an additional species-specific network using the *getKEGGModelForOrganism* function from RAVEN and the organism code corresponding to *Sus scrofa* (‘ssc’). The amino acid sequences for the genes in the species-specific network were retrieved from KEGG, and their subcellular localization for the genes in the species-specific network was predicted using DeepLoc 2.1 (Ødum et al., 2024). The results were used to assign compartments to the species-specific reactions and metabolites. Next, the species-specific network was added to the ortholog model using the *addMetabolicNetwork* function from Wang (Wang et al., 2018), and gene, reaction and metabolite IDs were edited to match the format of the ortholog model. Following automated reconstruction efforts, the model underwent extensive manual curation to resolve reaction directionality and compartment assignments, remove duplicated or unsupported reactions and fill metabolic gaps based on metabolic databases and literature. The resulting model was constrained based on media composition and experimentally determined uptake and excretion rates.

### Biomass Composition Measurements

Biomass composition measurements were performed on freshly harvested tissue samples provided by Prof. Robert Delmore from the Animal Sciences Department at Colorado State University. Muscle samples were taken from the semitendinosus muscle and adipose samples were taken from the pig back fat area. To obtain individual cells from the tissue samples for the biomass composition measurements, an adapted protocol of previously published cell isolation methods was used (Li et al., 2015). Samples were first sterilized with 70% ethanol in a petri dish and rinsed 4 times with cold PBS. Tissues were then cut into small pieces using sterile scissors under cold PBS in a plastic petri dish, transferred to a 50 mL conical tube and centrifuged at 1000*g* for 10 min. The supernatant was discarded and 3 volumes of collagenase digestion solution (Collagenase, Type II, concentration1.5 mg/mL dissolved in DMEM) were added. Tissues were incubated for 1 hour at 37°C. 15 mL of complete media (DMEM/Hams F12 (Gibco) supplemented with FBS (Fisherbrand)) was added to digested tissues and centrifuged at 2,000*g* for 10 min. Cells were resuspended in 15 mL of complete media and centrifuged at 2,000*g* for 10 min again. Cells were resuspended in 15 mL of complete media and successively filtered through 100 μm, 70 μm and 40 μm cell strainers. To determine cell dry weights, samples were lyophilized in a Buchi lyophilizer for 7 days to remove all liquid from the sample. After lyophilization, samples were dried in an 80°C oven overnight.

Biomass composition measurements were performed following previously published protocols (Meagher et al., 2024). To determine total protein content, 15 mg samples were resuspended in 0.2N NaOH and heated to boiling temperature for 1 hour and the resulting protein solution was quantified using a Pierce BCA protein assay kit for a microplate reader. Total lipid content was determined using a modified version of a chloroform: methanol extraction method previously described (Breuer et al., 2013). Briefly, 35-100 mg samples were suspended in 4 mL of a 2:2.5 chloroform:methanol solution, vortexed, and 2.5 mL DI water was added. The samples were vortexed again and centrifuged at 500*g* for 3 minutes. The chloroform phase was removed using a glass Pasteur pipette and transferred to a glass vial. An additional 2 mL of chloroform were added to the samples in chloroform: methanol solution to re-extract lipids, the samples were vortexed and centrifuged at 500 x *g* for 3 minutes. The chloroform fraction was again removed using a glass Pasteur pipette and transferred to the glass vial containing the first extract. The glass vials containing the extracted lipids in the chloroform layer were allowed to evaporate overnight in a chemical fume hood and weighed after all solvent had evaporated. To determine total carbohydrates, 1.5-3 mg of samples were heated to 100°C in a 75% H_2_SO4 solution containing 2 mg/mL anthrone for 15 minutes. A set of prepared glycogen standards (0, 40, 100, 200, and 400 mg/L) was treated in the same manner as the samples to create a standard curve. After incubating for 15 min at 100°C, the samples were immediately placed on ice and allowed to cool. After cooling to room temperature, a microplate reader was used to measure sample absorbance at 578 nm. A standard curve was constructed, and the carbohydrate content concentration was calculated. The nucleic acid content of the sample was determined using the Qubit double-stranded DNA and RNA broad range (BR) assay kits and the ATGC content based on the genome and transcriptome respectively. Detailed information on reagent brands and catalog numbers is provided in the Supplemental Information.

### Cell culture

Pig Muscle Satellite cells (pMSC) from Duroc pigs were first cultured in the media provided by the supplier and later cultured in DMEM/F12 (Gibco) supplemented with Skeletal Muscle Cell Growth Supplement (ScienCell), 5% FBS (Fisherbrand), and 1 mL Primocin (InvivoGen). pMSCs were cultured in cell culture flasks pre-coated with poly-l-lysine (MilliporeSigma) and grown at 37°C and 5% CO_2_. Cell culture media was changed every 24 hours, and cells were passaged using Trypsin-EDTA (0.25%) (Gibco) after reaching 70% confluency. Cells were not used for more than 6 passages. Detailed information on reagent brands and catalog numbers is provided in Table S1 in the Supplemental Information.

### Spent Media Analysis

Cells were seeded in 24-well plates, at a concentration of 1 x10^4^ cells/mL per well and cultured at 37 °C and 5% CO_2_. Four biological replicates were created for every time point and cell culture media was changed every 24 hours, depending on the experiment. At every given time point, the supernatant was removed from four wells and centrifuged to remove any cell and debris, and this portion was used for spent media analysis. Subsequently, 0.5 mL of Trypsin 0.25 % was added to the wells to remove the cells and count them. The cells were incubated at 37°C and 5% CO_2_ for 3-4 min until the cells lifted. Complete media was added to the well plates to neutralize the Trypsin reaction, cells were collected with cell scrapers, and the cells were centrifuged at 200 g for 5 min. The number of cells per well was counted manually or with a Scepter 3.0 Automated Cell Counter. The supernatant samples collected from the wells were analyzed using a Roche Cedex Bio Analyzer (glucose, glutamine, glutamate, lactate, pyruvate, and ammonia assays). The amino acid concentrations in the spent media were quantified using Gas Chromatography/Mass Spectrometry (GC/MS) (Antoniewicz et al., 2007) using 5 µL of Norvaline at 5 µmol/mL as the internal standard.

### Amino Acid Supplementation Simulations

Parsimonius Flux Balance Analysis (pFBA) was carried out on the genome-scale metabolic model with the muscle biomass formation equation used as the objective function to maximize biomass. We held the glucose uptake rate and all but one amino acid flux constant (based on experimental data) and then iterative increased the uptake flux of a single amino acid while recording the predicting growth rate. Constraints for these simulations are show in Table 1, and uptake rates were artificially increased by 0.01 mmol·gDW^-1^·hr^-1^ to simulate increased amino acid uptake.

**Table 1.**
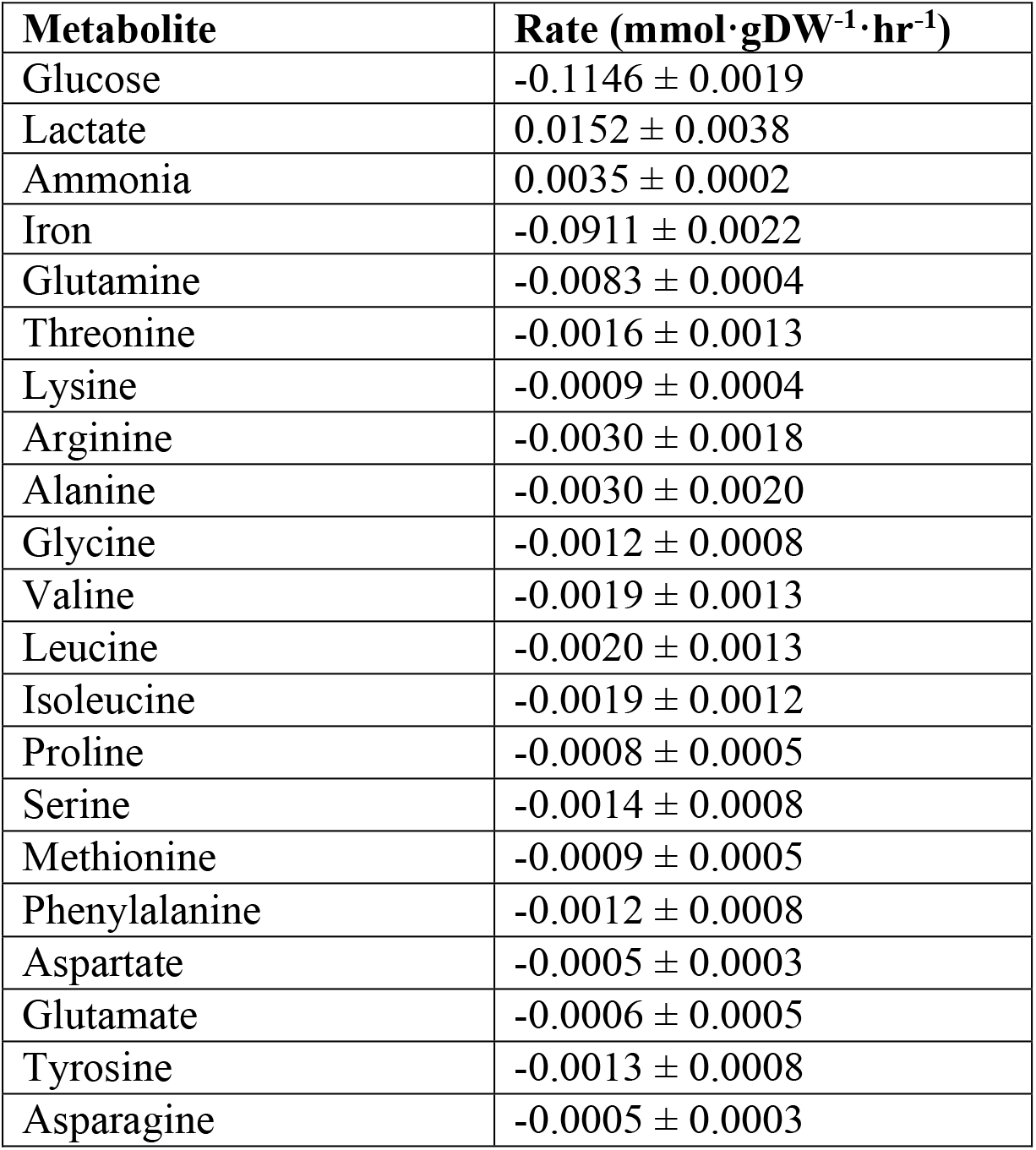
Experimentally determined exchange rates for dSMCs grown in DMEM/F12. Positive values indicate the metabolite is produced during growth, negative values indicate the metabolite is taken up by cells during growth. The reported error is theReplication Statement according to author guidelines standard deviation for n = 4 biological replicates.

### Amino Acid Supplementation Experiments

We selected the top seven performing amino acids for increased cell growth and increased them 4 fold from the basal DMEM/F12 concentrations. Cells were seeded in 24-well plates treated pre-coated with poly-l-lysine with 1 mL of cell mix at a concentration of 5 × 10^4^ cells/mL per well for all controls and treatment conditions. Cells were cultured in DMEM/F12 for the first 12 hours at 37 °C and 5% CO_2_ to allow for adhesion. After 12 hours, the cell media was changed to either control or treatment media (4x concentration of an individual amino acid). Cell culture media was changed every 24 hours. Cell samples and spent media analysis samples were taken at multiple timepoints (0, 6, 12, 24, 30, 36, 42, 48, 54, 72, 96, and 120 hours). At each time point, the supernatant media was removed, 0.5 mL of Trypsin 0.25% were added to the corresponding wells, and the cells were incubated at 37°C and 5% CO_2_ for 3 – 4 minutes until the cells lifted. Complete media (media supplemented with FBS, growth factors, and antibiotics) was added to the well plates to neutralize the Trypsin solution, the plate surface was scraped with cell scrapers, the cells were collected and counted using an Improved Neubauer Counting Chamber (Hausser Scientific) and Scepter Automated Cell Counter (Millipore). Growth rates and doubling times were calculated using equations 1 and 2 below (Saad et al., 2023):

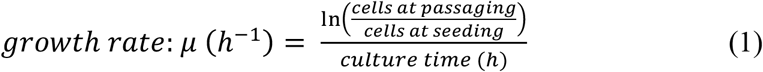

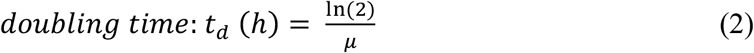

### Statistical Analysis

To compare more than two samples, a one-way Analysis of Variance (ANOVA) was performed, along with Dunnett’s test to test the significance of multiple treatment groups compared to the control group. An additional Kruskal-Wallis test was performed on the growth rate and doubling experimental results. A Kruskal-Wallis test a non-parametric test and can be described as a one-way ANOVA on rank. *P* values < 0.05 were treated as significant. All reported errors represent the standard deviation of the data.

## RESULTS

### Model reconstruction

The final manually curated model, iSsus3744 (Supplemental File 1) consists of 3,744 genes, 8,854 metabolites and 12,248 reactions in 8 subcellular compartments (See Figure 1). The full list of reactions and metabolites in the model are provided in Supplemental Files 2 and 3 respectively. The model has a MEMOTE score of 78%, indicating a consistent model with genomic support and complete annotations (Christian Lieven et al., 2020). More detailed information on the MEMOTE score of this genome-scale model can be found in Supplemental File 4.

**Figure 1.**
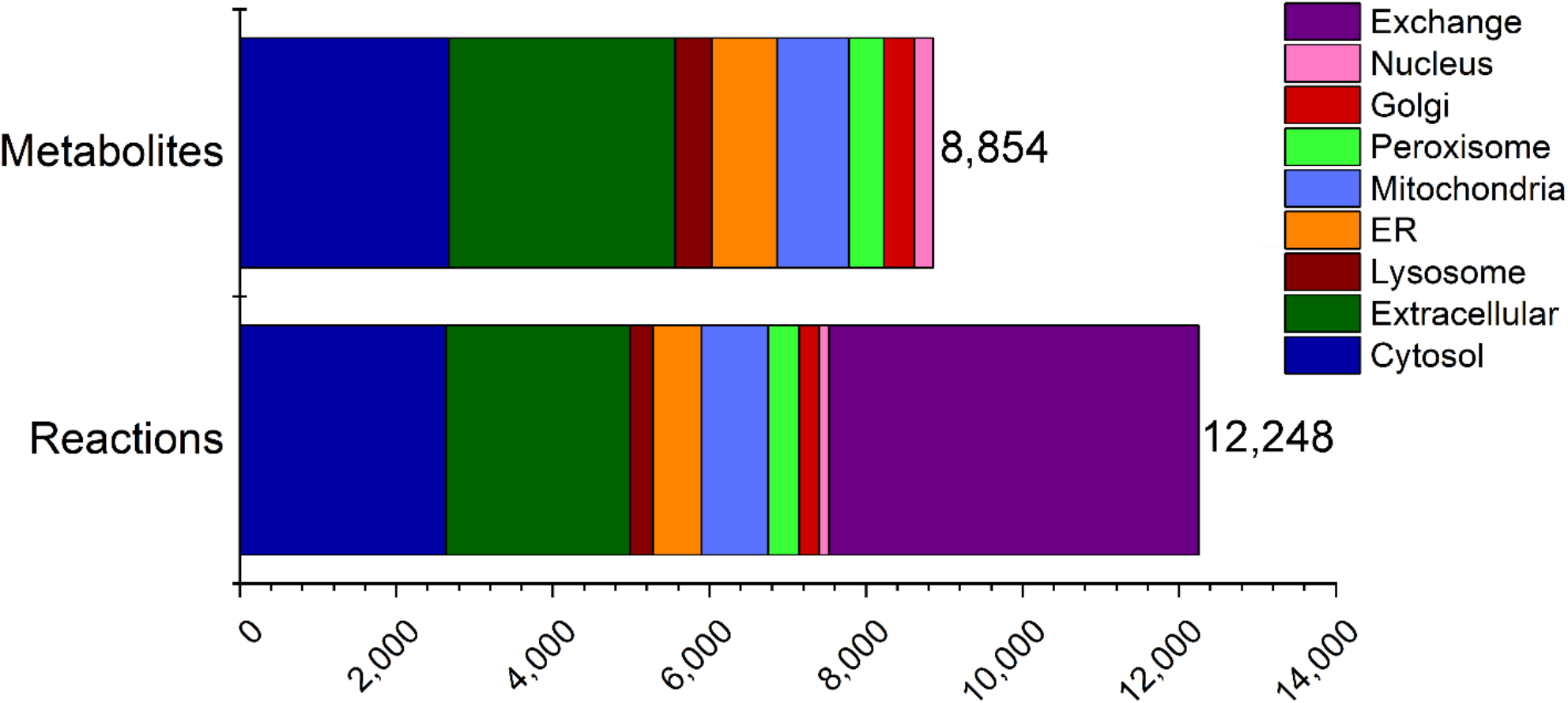
iSsus3744 model summary. The pig genome scale model represented 3744 genes, 8,854 metabolites and 12,248 reactions in 8 cellular compartments. ER – Endoplasmic Reticulum.

### Experimentally Measured Biomass and Model Constraints

Biomass composition was measured in two types of porcine tissue: adipose and muscle in order to construct an experimentally determined biomass formation equation for the pFBA model. The macromolecular composition of each tissue was measured to determine total carbohydrate, lipid, protein, and nucleotide content. Additional analyses were performed to measure the composition of each macromolecule in terms of its associated monomers, i.e. amino acids, sugars, and fatty acids (see Figure 2). Our results show that adipose tissue has 88.5% more relative arginine content, 57.3% more relative glycine content, and 17.8% more alanine content than muscle tissue. Muscle tissue had a higher composition of leucine, isoleucine, proline, serine, threonine, methionine, phenylalanine, asparagine, glutamate, lysine, and tyrosine compared to fatty tissue. Of these, asparagine, leucine, and isoleucine had a more marked difference compared to fatty tissue. Muscle tissue has 79.8% more relative content of asparagine, 70.7% more relative content of isoleucine, and 63.8% more relative content of leucine than fatty tissue. Fatty tissue has 96.7% more relative composition of oleic acid (cis-9 oleic acid), 49.4% more relative content of palmitate, 86.4% more relative content of octadecenoate (stearate), and 68.3% more relative content of trans 9,12-octadecanoate (linolelaidate) compared to muscle tissue. Muscle tissue had 97.3% more relative content of Eicosatetraenoic acid (ETA) (cis-5,8,11,14 eicosatetraenoic acid), and 66.1% more relative content of myristate than fatty tissue. A table of the biomass formation equation for each tissue type is provided as Table S2 in the Supplemental Information.

**Figure 2.**
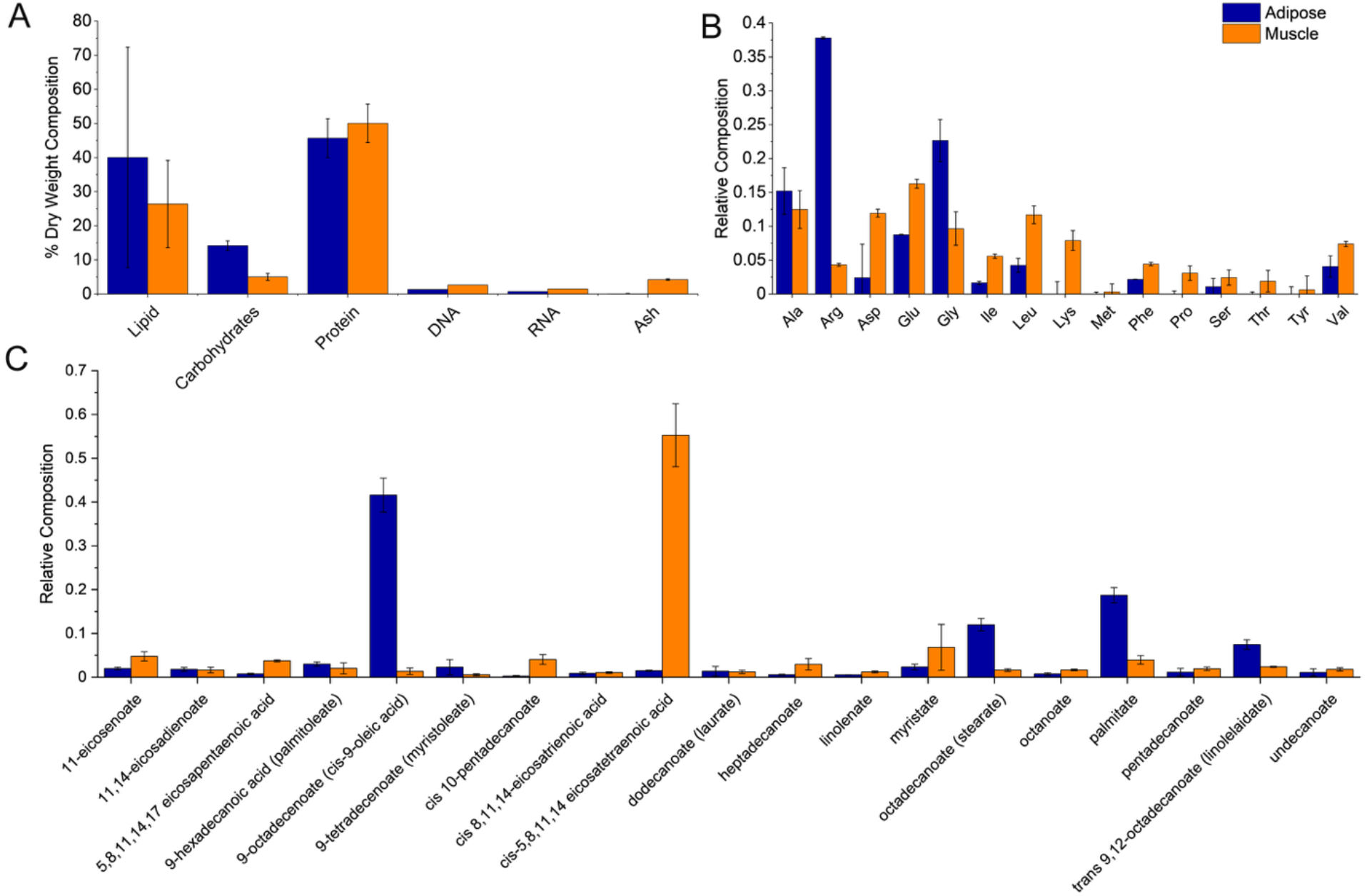
Average biomass composition measurements of two *S. scrofa* tissue types: adipose and muscle. A) Biomass composition in terms of major macromolecules, B) amino acid composition of hydrolyzed proteins and C) fatty acid composition. Error bars represent standard deviation of n = 3 samples.

### Flux Variability Analysis

Flux variability analysis (FVA) was performed to determine the allowable range of fluxes for substrate and product exchanges that still satisfied the constraints on the model (see Figure 3). Essential amino acids (Ile, Leu, Lys, Met, Phe, Thr, Val) are required to be taken up from the medium because the model doesn’t have the reactions necessary to synthesize these amino acids on its own. The remaining amino acids all have much larger ranges of allowable flux spanning both uptake and excretion. If the ranges of allowable fluxes are sorted in order of size (smallest to largest), then those with the narrowest acceptable flux ranges are implied to be those most critical to growth. In order of importance based on the metabolic model, this would be Met, Arg, Val, Leu, Lys, Ile, Phe, Tyr, Thr, Asn, Asp, Pro, Glu, Gln, Gly, Ser, Ala. Ammonia and lactate are both typical byproducts of cell growth that negatively impact cell growth when they accumulate (Hassell et al., 1991). The FVA analysis indicates that, at least stoichiometrically, these fluxes can be adjusted by altering the uptake of other components of the media and thus minimize their impact on growth.

**Figure 3.**
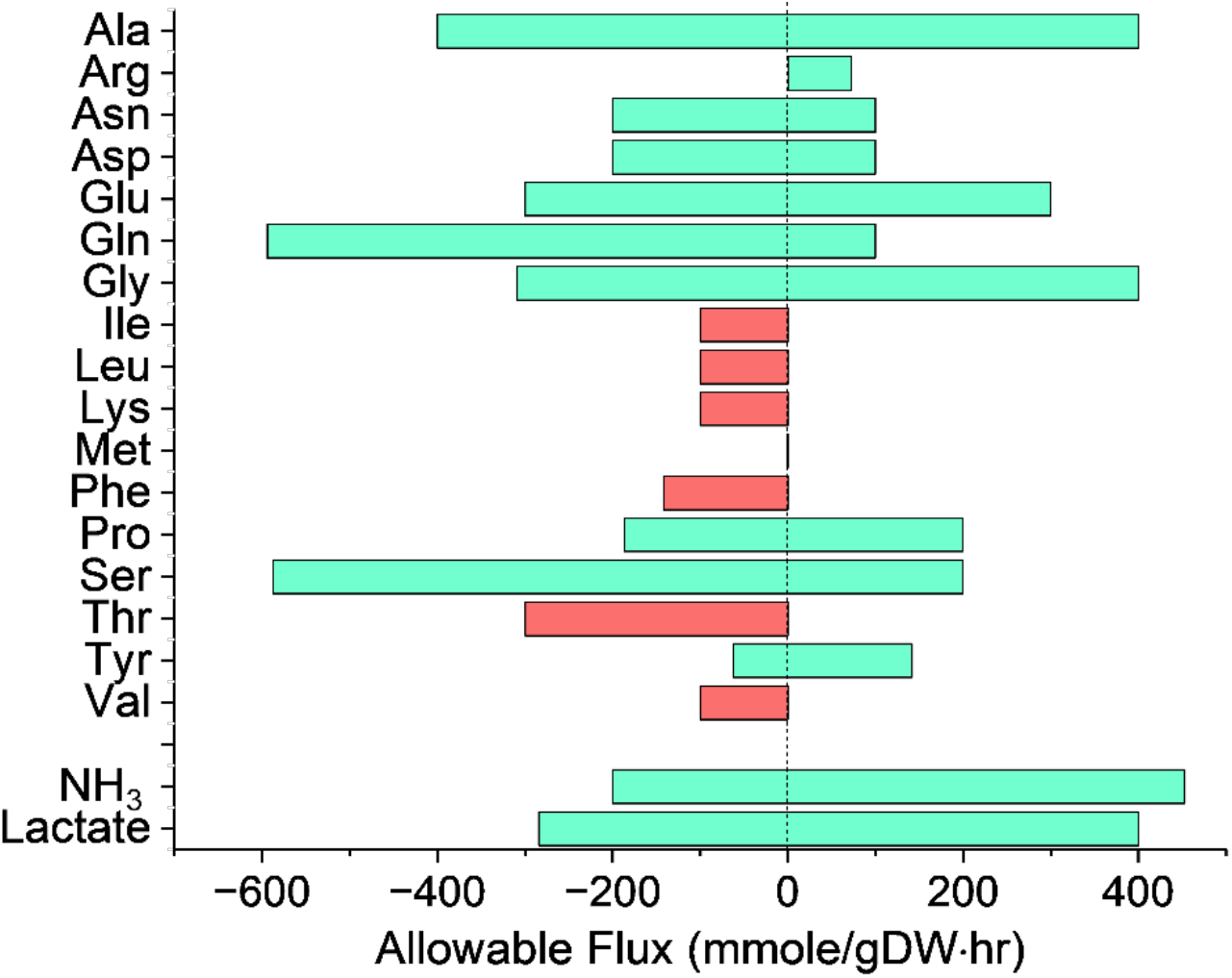
Flux variability analysis for uptake and excretion rates for dMSCs. FVA (Flux Variability Analysis) is a tool for quantifying the feasible ranges of reaction fluxes in a metabolic network at optimal production, allowing analysis of the flexibility of reactions within the network. FVA here is shown with the objective of maximizing biomass. Our results show that the model has a wider range of feasibility in reaction fluxes of alanine, glutamine, glycine, ammonia and lactate. Isoleucine, leucine, methionine, phenylalanine, threonine, and valine have a reduced range of feasibility, meaning changes in those uptake rates have more of an impact on growth rates.

### Media Optimization Simulations

The genome scale model was used to identify changes in culture media that can lead to improved growth. One of the most challenging aspects of mammalian cell culture is the accumulation of metabolic byproducts, such as lactate and ammonia (Martins et al., 2024), which are toxic to the cell and impede growth. pFBA was used to investigate if cells would be capable of growth at the same rate if we forced the model to minimize the production of either of these products. The model simulations illustrate that it is *stoichiometrically* feasible to grow at the same growth rate without producing either byproduct. We examined the exchange fluxes for both cases (see Figure 4) to determine how the demand for amino acids changed in each case. To minimize the production of lactate, the model shows a decrease in the uptake of aspartate glutamine, leucine, lysine, threonine and valine. The essential amino acids have a non-zero uptake flux, but the non-essential amino acids of aspartate and glutamine drop to zero. The minimize lactate simulation also shows a dramatic decrease in the production of asparagine. In reducing lactate production to zero, the ammonia production does increase 1.5 fold over the control (growth in DMEM/F12). The simulation to minimize the production of ammonia also predicts a decrease in glutamine, but an increase for every other amino acid.

**Figure 4.**
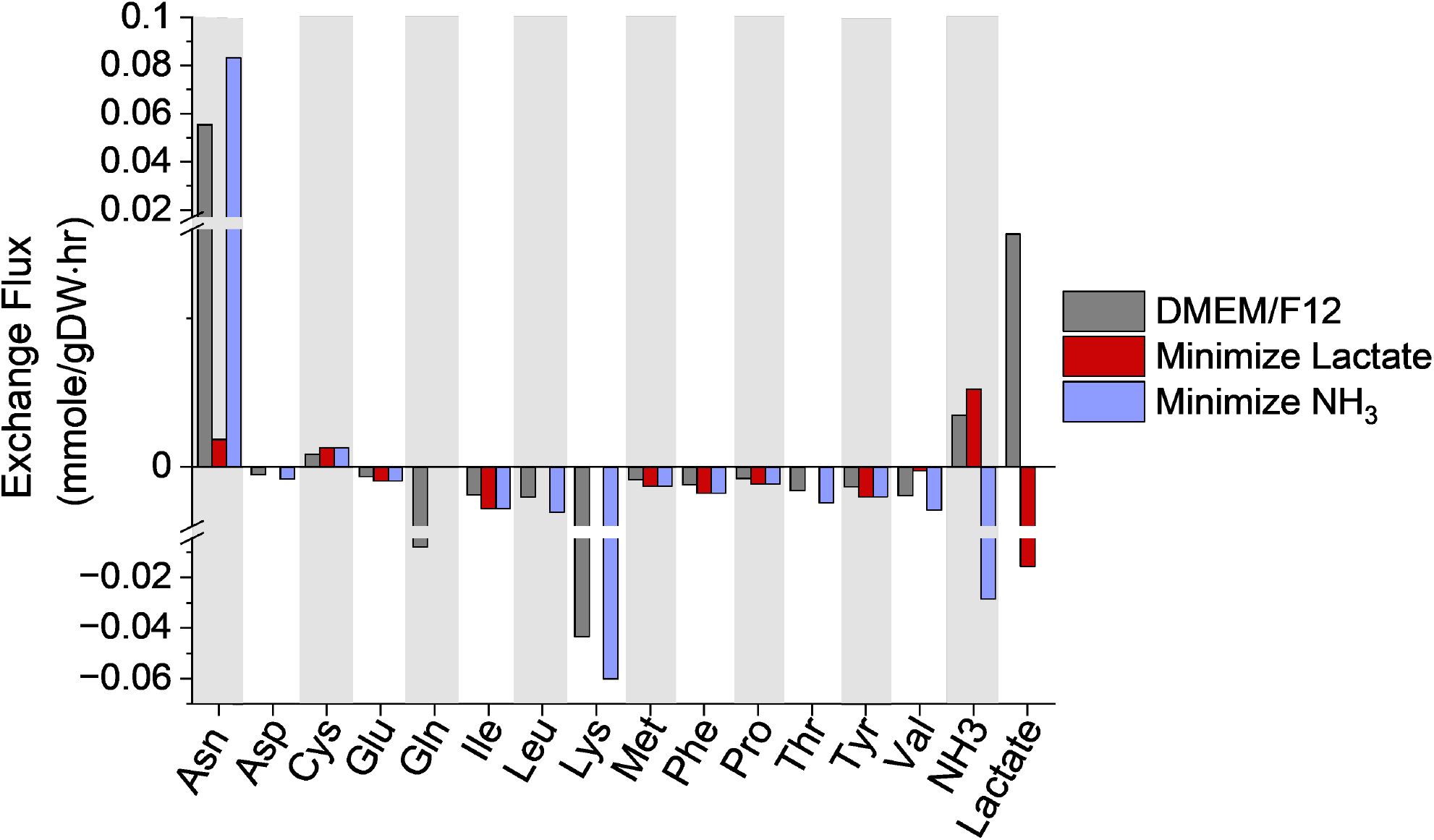
pFBA simulations to maximize biomass and minimize production of toxic byproducts (NH_3_ and Lactate).

Our porcine muscle satellite cell line was grown in DMEM/F12 medium, a widely used mammalian cell culture growth medium. In an effort to tune this basic formulation for our dMSC cell line, pFBA was used to identify amino acids whose supplementation would have the largest positive impact on growth. In brief, we incrementally increased the uptake rate of each amino acid one at a time, ran the simulation to optimize biomass and recorded the predicted growth rate. The amino acids that lead to the highest increases in biomass, shown in Figure 5 with the steepest slopes 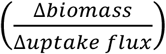, were isoleucine, leucine, methionine, phenylalanine, proline, tyrosine and valine. The amino acids that are not included in **Error! Reference source not found**. have little to no impact on growth; the complete data set is given in Supplemental File 5.

**Figure 5.**
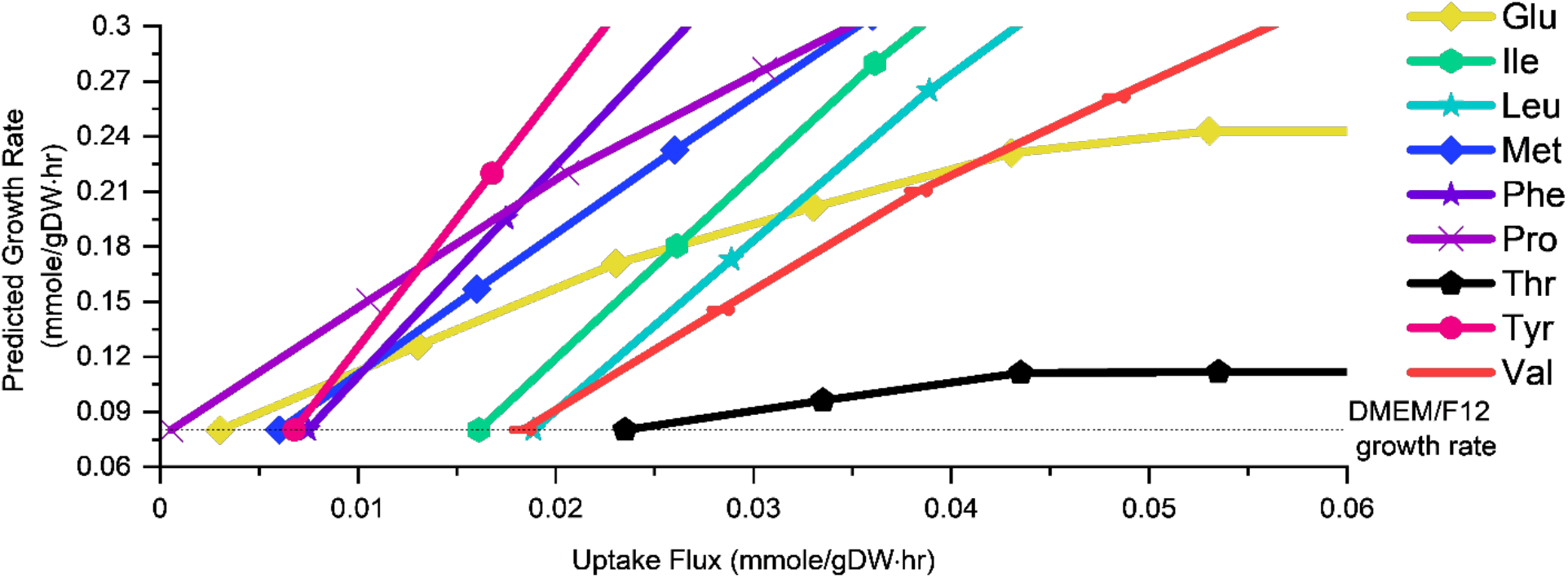
pFBA simulations of amino acid supplementation. Simulations were performed for all 20 amino acids, with the top performing amino acids shown. To carry out these simulations, the uptake rate of glucose was held constant while the uptake rate of each amino acid was increased individually. The simulated growth rate was plotted as a function of the uptake rate of the variable amino acid. Based on these simulations, we chose phenylalanine, leucine, isoleucine, tyrosine, valine and lysine to carry out the amino acid supplementation experiment.

### Experimental Validation of Model Predictions

To test the accuracy of model predictions, we grew the dMSCs in DMEM/F12 supplemented a 4-fold increase of each of the amino acids that were predicted to have a positive impact on growth (isoleucine, leucine, methionine, lysine, phenylalanine, tyrosine, and valine), the results of the experiment are given in Figure 6. The control (DMEM/F12 medium) had a growth rate of 0.022 ± 0.002 (h^-1^), which translates to a doubling time of 31.9 hours. Most of the treatment groups had higher average growth rates but were not statistically significant when we used Dunnett’s test to compare across treatments groups. The supplementation of one amino acid, phenylalanine was determined to be statistically significant. The growth rate for cells supplemented with Phe was μ = 0.04 ± 0.008 (h^-1^), which translates to a doubling time of 17.2 hours, a 46% reduction in doubling time. This is a significant reduction given that only a single amino acid concentration was changed; it also indicates that significant improvements in cell culture can be achieved by optimizing media using this technique.

**Figure 6.**
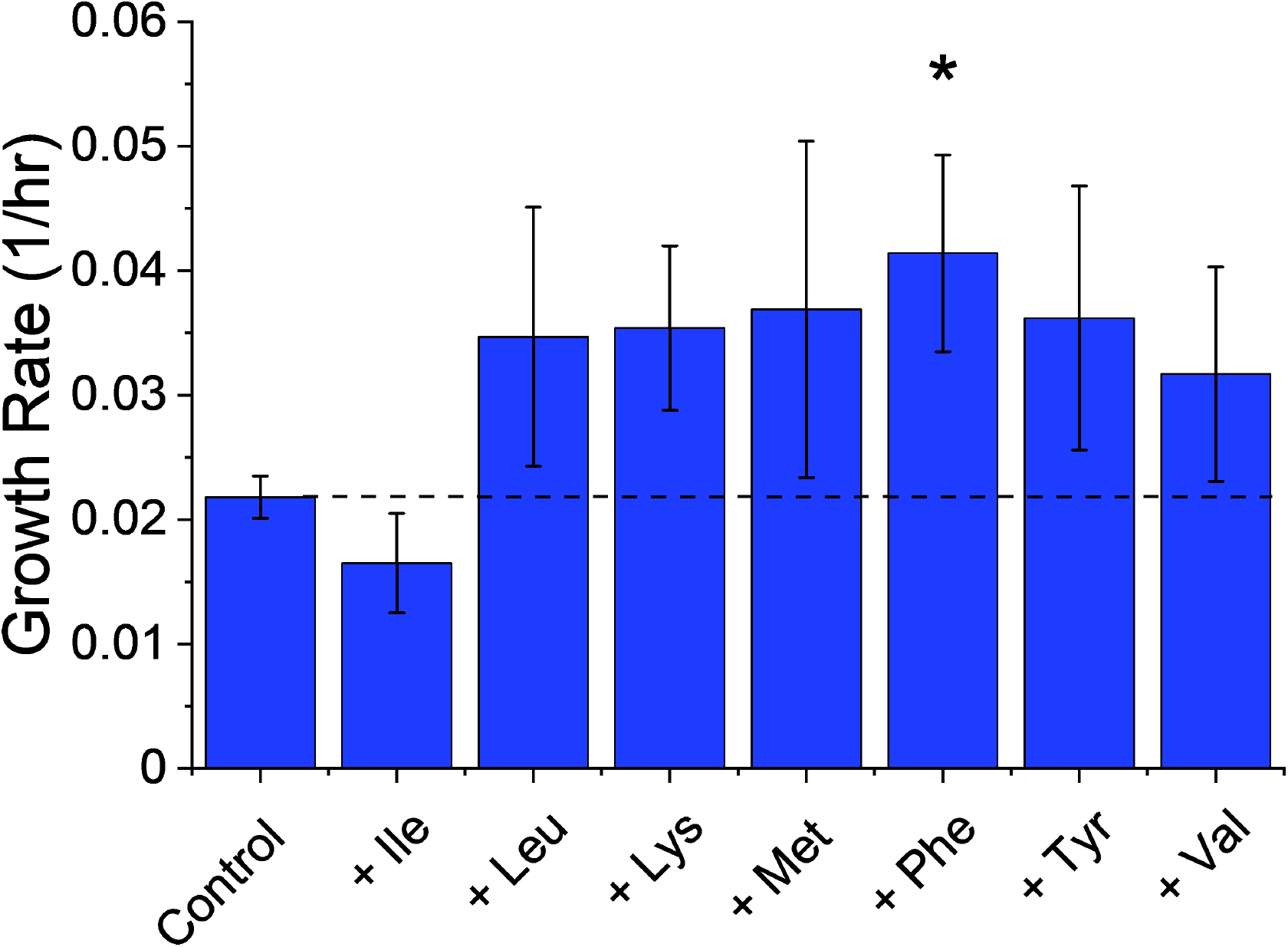
Experimentally measured growth rates for dSMCs grown in the control medium (DMEM/F12) and with 4x supplementation of the indicated amino acids. A Dunnet’s test of the impact on growth indicates that supplementation of phenylalanine is the only statistically significant change compared to all treatments. Error bars represent standard deviation of *n* = 4 samples.

## DISCUSSION

### Model Development

Genome-scale metabolic models (GEMs) remain relatively underdeveloped within the field of cellular agriculture. To date, GEMs have been constructed for only a limited number of agriculturally relevant species: cattle (Junkyu Lee, 2024; Kim et al., 2016), chicken (Salehabadi et al., 2022), shrimp (Gao et al., 2021) and salmon (Zakhartsev et al., 2022). A small metabolic network with 97 reactions has also been reported for duck (Lohr et al., 2014). Unfortunately, the quality and predictive performance of these models varies considerably, largely reflecting limited experimental datasets and a lack of rigorous validation. One standard measure of model quality is the MEMOTE (METabolic Model Tests) score (C. Lieven et al., 2020), which uses an automated algorithm to calculate a score for genome-scale metabolic models based on 4 general areas: annotation, basic tests, biomass reaction and stoichiometry and higher scores typically indicate a higher quality model. The most recent genome-scale model for cattle has a MEMOTE score of 79% (Junkyu Lee, 2024; Kim et al., 2016), and salmon (SALARECON) is 96% (Zakhartsev et al., 2022); neither the chicken or shrimp models report MEMOTE scores and are not provided in the proper format to run MEMOTE. The cattle model comprises 2,986 genes, 13,278 reactions and 8,652 metabolites, a similar size to the pig model we report here and has a similar MEMOTE score. The salmon model is significantly smaller, with 718 reactions and 530 metabolites and was constructed manually which contributes to the higher MEMOTE score.

While MEMOTE provides a standard checklist for assessing genome-scale metabolic models, relying on it as a primary indicator of model quality is problematic. Its scoring system emphasizes structural features such as annotation completeness, mass and charge balance, and network consistency rather than the model’s predictive accuracy or biological validity. As a result, a model can achieve a high MEMOTE score yet still perform poorly in simulating experimentally observed phenotypes or fluxes. For example, the iSsus3744 MEMOTE score is lower because the MEMOTE algorithm only searches for references to a limited number of annotation databases, especially for gene annotations. Although this model has a score of 100% for gene annotation, the gene score subtotal being added to the total score for gene is only 33% because MEMOTE does not search for Ensembl gene annotations. Additionally, MEMOTE does not adequately capture context-specific performance, such as condition-dependent behavior, parameterization (e.g., maintenance energy), or the integration of experimental constraints or omics data, which are often critical for real-world applications. The iSsus3744 model has the additional benefit of experimentally determined constraints for uptake and excretion fluxes as well as experimentally determined biomass composition data. It is important to note that experimental data is essential for constraining, validating and ensuring the predictive reliability of genome-scale metabolic models and that this is beyond what standardized scoring metrics, such as MEMOTE, can capture.

### Improving media formulation for porcine cell culture

Our pMSC cell line grown in DMEM/F12 media supplemented with FBS had a doubling time of 31.9 ± 2.6 hours, which agrees with reported doubling times of primary muscle satellite cells in the range of 18.6 ± 0.8 h to 51.36 hours (Yang et al., 2017; Zhu et al., 2013). Primary cell lines isolated from neonatal pigs are reported to have increasing doubling time with increasing passage, from 32.81 hours in passage 2 to 51.36 hours in passage 10 (Yang et al., 2017). The age of the animal when primary cells are harvested also impacts growth rate; cells harvested from young animals have a faster doubling time than adult animals (Zhu et al., 2013). For all the amino acid supplementation experiments, the same primary muscle satellite cell line was used to minimize differences due to the age of the animal or passage number.

Based on the results of our amino acid supplementation model simulations, we investigated how the supplementation of 7 individual amino acids (Ile, Leu, Lys, Met, Phe, Tyr and Val) impacted the growth rate of our pMSC cell line. Isoleucine was predicted by the model to increase growth but had the opposite effect in our experiment. Animal studies have reported that in low protein diets, supplementation of Ile has either no impact or negative impact on feed efficiency; whereas Ile supplemented with Val improves feed intake and feed efficiency (Nørgaard & Fernández, 2009). The addition of phenylalanine increased growth rate by 89%, the addition of lysine increased growth rate by 62%, and the addition of tyrosine increased growth rate by 65%. Methionine and leucine also improved growth rates by 69% and 59%, respectively (see supplemental information Table S3). A one-way ANOVA analysis indicates that there is indeed a statistical difference amongst the group; a Dunnett’s test indicates that supplementation of phenylalanine id the only statistically significant change in growth. Animal science studies on pig metabolism have shown that these amino acids are important for the growth and development of piglets. The limiting amino acids in piglet diets in order of importance are lysine, methionine, threonine, tryptophan, valine, isoleucine and leucine, phenylalanine, and histidine (van Milgen & Dourmad, 2015). Phenylalanine and tyrosine are important amino acids for the growth of pigs; the estimated requirement range is 57-61% of standardized ileal digestible relative to lysine (Gloaguen et al., 2014). Phenylalanine and tyrosine metabolism are deeply connected, and phenylalanine can be converted to tyrosine through an irreversible hydroxylation (Gloaguen et al., 2014; Matthews, 2007). Improved growth rate after addition of lysine is expected, given that lysine is the main limiting amino acid in swine diets and has been reported to impact growth performance (Camp Montoro et al., 2022; Hasan et al., 2020; Kwon et al., 2022). The addition of lysine at a ratio of 1.45% of lysine/amino acid fraction has been shown to increase growth and weight gain in slowly growing pigs (Camp Montoro et al., 2022). Methionine is the second or third limiting amino acid in pig diets (depending on the stage of life of the pig) and plays an important role in promoting development and growth in piglets (Wan et al., 2025). These findings suggest that future media optimization efforts for cell lines derived from agriculturally relevant species could benefit from animal science studies and dietary guidelines for the animal from which the cells line are derived. Given the wealth of data for optimizing growth in pigs also suggests that future efforts to optimize cell culture media should focus on balancing the ratio of amino acids.

### Genome scale models in cellular agriculture

The use of genome-scale metabolic model-guided media design represents a powerful and innovative strategy for streamlining cell culture media development and optimization. In this work, we demonstrate that a metabolic model can successfully identify limiting amino acids and accurately predict their effects on cell growth, highlighting the practical value of systems-level modeling approaches for rational media design.

FBA provides an efficient framework for predicting optimal metabolic states and maximizing objectives such as growth or metabolic production. While FBA can yield multiple feasible solutions (Park et al., 2010), the incorporation of experimentally determined constraints, as performed in this study, substantially improves prediction accuracy and biological relevance. Additionally, although traditional FBA assumes steady-state metabolism and is therefore best suited for defined metabolic windows, advanced approaches such as dynamic FBA (dFBA), extend these capabilities by incorporating temporal metabolic changes and enabling the study of dynamic cellular behavior. Integration of genetic constraints to restrict the availability of metabolic reactions using RNA-seq data, such as was done to model dynamic changes in metabolism due to shifting light in algae (Metcalf & Boyle, 2022), can also increase accuracy of predictions. Flux variability analysis (FVA) is another tool that offers valuable insight into the flexibility and robustness of metabolic networks by quantifying allowable flux ranges under specific constraints. Metabolic models provide a strong computational foundation for interrogating metabolism and identifying key pathways influencing ellular performance. Importantly, these approaches also serve as a basis for the development of increasingly sophisticated models that integrate additional layers of biological regulations, including transcriptional regulations, signaling pathways and growth factor responses.

## CONCLUSION

In this study, we developed iSsus3744, the first genome-scale metabolic model for *Sus scrofa*, and demonstrated its utility as a predictive platform for rational media optimization in cultivated pork production. By integrating experimentally determined biomass composition and extracellular flux measurements, we generated a biologically constrained model capable of identifying amino acid limitations that restrict cell proliferation. Experimental validation confirmed that model-guided supplementation strategies significantly improved the growth of porcine muscle satellite cells, with phenylalanine supplementation reducing doubling time by nearly half. This work highlights the value of combining systems biology and experimental validation to accelerate media optimization for cultivated meat applications. Beyond identifying limiting nutrients, iSsus3744 establishes a framework for integrating additional biological complexity, including transcriptional regulation, signaling pathways, and dynamic metabolic behavior. The model also provides a foundation for future isotope-assisted metabolic flux analysis studies and the development of serum-free, cost-effective culture media. Collectively, these findings demonstrate that genome-scale metabolic modeling can play a transformative role in advancing cultivated meat bioprocess development and improving our understanding of porcine cell metabolism.

## Acknowledgments

We would like to thank Prof. Robert Delmore from the Department of Animal Sciences at Colorado State University for providing tissue samples for biomass composition studies. We would like to thank the Cultivated Meat Modeling Consortium for the fruitful connections and funding that partially supported this research. And we would like to thank Elliot Swartz, from the Good Food Institute, for his mentorship and fruitful discussions during a Research Fellowship for SG and for sharing insights about the Cultivated Meat industry.

