## Supplemental Info for "iSsus3744: A Genome-Scale Model-Guided Strategy for Rational Media Design for Cultivated Pork"

### **SUPPLEMENTAL INFORMATION**

**Table S1**. Detailed information for reagents used for cell culture.

| **Reagent** | **Brand** | **Catalog Number** |
| --- | --- | --- |
| Collagenase Type II | Gibco™ | 17101015 |
| BenchStable DMEM/F12 | Gibco™ | A4192001 |
| Fetal Bovine Serum (FBS) | Fisherbrand^TM^ | FB12999102 |
| Pierce BCA protein assay kit | Thermo Fisher | 23225 |
| Anthrone reagent | Thermo Scientific Chemicals | 104961000 |
| Pig Muscle Satellite Cells | AcceGen | ABC-TC4019 Duroc |
| Muscle Satellite Cells Medium | AcceGen | ABM-TM4019 |
| Skeletal Muscle Cell Growth Supplement | ScienCell | 3552 |
| Adipocyte Growth Supplement | ScienCell | 7262 |
| Primocin | InvivoGen | NC9141851 |
| Poly-l-lysine | MilliporeSigma | A005C |
| Recombinant Murine FGF | ScienCell | 124-02 |
| HematoGro Supplement | ScienCell | 5552 |
| Trypsin-EDTA (0.25%) | Gibco™ | 25200072 |
| L-Serine (Cell Culture Reagent) | Thermo Scientific Chemicals | AAJ6218709 |
| L-Isoleucine (Cell Culture Reagent) | Thermo Scientific Chemicals | AAJ6304514 |
| L-Proline, Cell Culture Reagent) | MP Biomedicals | ICN19472825 |
| L-Glutamic Acid, 99+% Monosodium Salt | MP Biomedicals | ICN19467780 |
| L-Alanine (Cell Culture Reagent) | Thermo Scientific Chemicals | AAJ6027918 |
| L-Valine | Thermo Scientific Chemicals | AC140810250 |
| L-Tyrosine disodium salt dihydrate | Thermo Scientific Chemicals | AAJ6177022 |
| L-Tryptophan (Cell Culture Reagent) | Thermo Scientific Chemicals | AAJ6250809 |
| L-Threonine (Cell Culture Reagent) | Thermo Scientific Chemicals | AAJ6370930 |
| L-Glycine (Cell Culture Reagent) | Thermo Scientific Chemicals | AAJ6240722 |
| L-Leucine (Cell Culture Reagent) | Thermo Scientific Chemicals | AAJ6282422 |
| L-Methionine (Cell Culture Reagent) | Thermo Scientific Chemicals | AAJ6190418 |
| L-Phenylalanine (Cell Culture Reagent) | Thermo Scientific Chemicals | AAJ6392522 |
| L-Lysine Hydrochloride | MP Biomedicals | ICN19469780 |
| L-Histidine monohydrochloride monohydrate | Thermo Scientific Chemicals | AAA1762718 |
| L-Cystine dihydrochloride, 99% | Thermo Scientific Chemicals | J62292.14 |
| L-Arginine Hydrochloride | Fisher BioReagents | BP372-100 |
| L-glutamine (Gibco GlutaMAX) | Gibco™ | 35-050-061 |

**Table S2**. Biomass formation equation stoichiometry for Duroc porcine muscle satellite cells to produce X g DW of biomass.

| **Metabolite ID** | **Metabolite Name** | **Coefficient** |
| --- | --- | --- |
| M_SUS00017_c | (13Z)-eicosenoic acid | -5.03E-05 |
| M_SUS00117_c | (7Z)-tetradecenoic acid | -8.32E-06 |
| M_SUS01362_c | arachidonate | -5.96E-04 |
| M_SUS01721_n | DNA | -2.63E-02 |
| M_SUS02006_c | glycyl-tRNA(gly) | -9.08E-04 |
| M_SUS02335_c | L-alanyl-tRNA(ala) | -9.41E-04 |
| M_SUS02340_c | L-arginyl-tRNA(arg) | -1.48E-04 |
| M_SUS02342_c | L-aspartyl-tRNA(asp) | -5.61E-04 |
| M_SUS02344_c | lauric acid | -1.99E-05 |
| M_SUS02377_c | L-glutamyl-tRNA(glu) | -4.01E-04 |
| M_SUS02387_c | linoleate | -2.76E-05 |
| M_SUS02401_c | L-isoleucyl-tRNA(ile) | -2.64E-04 |
| M_SUS02404_c | L-leucyl-tRNA(leu) | -5.54E-04 |
| M_SUS02405_c | L-lysyl-tRNA(lys) | -3.30E-04 |
| M_SUS02408_c | L-methionyl-tRNA(met) | -1.30E-05 |
| M_SUS02412_c | L-phenylalanyl-tRNA(phe) | -1.61E-04 |
| M_SUS02415_c | L-prolyl-tRNA(pro) | -1.70E-04 |
| M_SUS02416_c | L-seryl-tRNA(ser) | -1.49E-04 |
| M_SUS02419_c | L-threonyl-tRNA(thr) | -1.00E-04 |
| M_SUS02421_c | L-tyrosyl-tRNA(tyr) | -2.05E-05 |
| M_SUS02423_c | L-valyl-tRNA(val) | -3.99E-04 |
| M_SUS02456_c | margaric acid | -3.59E-05 |
| M_SUS02494_c | myristic acid | -9.84E-05 |
| M_SUS02642_c | octanoic acid | -3.78E-05 |
| M_SUS02646_c | oleate | -1.58E-05 |
| M_SUS02674_c | palmitate | -5.05E-05 |
| M_SUS02675_c | palmitolate | -2.60E-05 |
| M_SUS02690_c | pentadecylic acid | -2.62E-05 |
| M_SUS02938_c | stearate | -1.88E-05 |
| M_SUS03063_c | tRNA(ala) | -9.41E-04 |
| M_SUS03064_c | tRNA(arg) | -1.48E-04 |
| M_SUS03066_c | tRNA(asp) | -5.61E-04 |
| M_SUS03069_c | tRNA(glu) | -4.01E-04 |
| M_SUS03070_c | tRNA(gly) | -9.08E-04 |
| M_SUS03072_c | tRNA(ile) | -2.64E-04 |
| M_SUS03073_c | tRNA(leu) | -5.54E-04 |
| M_SUS03074_c | tRNA(lys) | -3.30E-04 |
| M_SUS03075_c | tRNA(met) | -1.30E-05 |
| M_SUS03076_c | tRNA(phe) | -1.61E-04 |
| M_SUS03077_c | tRNA(pro) | -1.70E-04 |
| M_SUS03078_c | tRNA(ser) | -1.49E-04 |
| M_SUS03079_c | tRNA(thr) | -1.00E-04 |
| M_SUS03081_c | tRNA(tyr) | -2.05E-05 |
| M_SUS03082_c | tRNA(val) | -3.99E-04 |
| M_SUS03117_c | undecylic acid | -3.15E-05 |
| M_SUS03161_c | glycogen | -5.01E-02 |
| M_SUS20165_u | (8Z,11Z,14Z)-Eicosatrienoic Acid | -1.15E-05 |
| M_SUS20578_u | Cis,Cis-11,14-Eicosadienoic Acid | -1.74E-05 |
| M_SUS21291_u | 5,8,11,14,17-Eicosapentenoic Acid | -4.07E-05 |

**Table S3. Experimentally determined growth rates for dSMCs grown in DMEM/F12 with amino acid supplementation.** The reported error is the standard deviation for n = 4 biological replicates.

| **Group** | **Growth rate (h^-1^)** | **Doubling time (h)** |
| --- | --- | --- |
| Control (DMEM/F12) | 0.022 ± 0.002 | 31.9 ± 2.6 |
| + 4x Phe | 0.041 ± 0.008 | 17.2 ± 3.4 |
| + 4x Lys | 0.035 ± 0.007 | 20.1 ± 3.7 |
| + 4x Tyr | 0.036 ± 0.011 | 20.4 ± 5.5 |
| + 4x Met | 0.037 ± 0.022 | 24.3 ± 13.2 |
| + 4x Leu | 0.035 ± 0.01 | 21.8 ± 8.3 |
| + 4x Val | 0.032 ± 0.009 | 23.1 ± 5.8 |
| + 4x Ile | 0.017 ± 0.004 | 43.7 ± 9.5 |
